# A redox-regulated type III metacaspase controls cell death in a marine diatom

**DOI:** 10.1101/444109

**Authors:** Shiri Graff van Creveld, Shifra Ben-Dor, Avia Mizrachi, Uria Alcolombri, Amanda Hopes, Thomas Mock, Shilo Rosenwasser, Assaf Vardi

## Abstract

Programmed cell death (PCD) in marine phytoplankton was suggested as one of the mechanisms that facilitates large scale bloom demise. Yet, the molecular basis for algal PCD machinery is rudimentary. Metacaspases are considered ancestral proteases that regulate cell death, but their activity and role in algae are still elusive. Here we biochemically characterized a recombinant metacaspase 5 from the model diatom *Phaeodactylum tricornutum* (PtMC5), revealing calcium-dependent protease activity. This activity includes auto-processing and cleavage following arginine. PtMC5 overexpressing cells exhibited higher metacaspase activity and were more sensitive to a diatom-specific infochemical compared to WT cells. Mutagenesis of potential disulfide-forming cysteines decreased PtMC5 activity, suggesting redox regulation. This cysteine pair is widespread in diatom type III metacaspases, but was not found in any other taxa. The characterization of a cell death associated protein in marine phytoplankton will enable deeper understanding of the ecological significance of PCD in bloom dynamics.

---

Diatoms are responsible for about half of marine photosynthesis, and therefore play a significant role in global biogeochemical cycles and in carbon sequestration^1,2^. Their evolutionary and ecological success in contemporary oceans suggests that diatoms may possess unique mechanisms for adaptation to diverse environmental conditions^3^. Diatoms can form massive blooms that are controlled by the availability of inorganic nutrients and light, and by biotic interactions with grazers, bacteria and viruses^4–8^. During these biotic interactions, an array of bioactive compounds (infochemicals) are produced, some of which are involved in allelopathic interactions and can regulate cell fate, hence shaping population dynamics and species composition^9^. The discovery of programmed cell death (PCD) in unicellular organisms raises fundamental questions about the origin of these self-destructing pathways and how a lethal phenotypes can prevail in the evolution of a single celled microorganism^10^. For instance, when diatom populations are subjected to grazing or nutrient stress, cells can rapidly induce the biosynthesis of diatom-derived oxylipins such as (*E,E*)-2,4-Decadienal (DD), a bioactive signal that may act as a chemical defense mechanism against grazing^11–13^. In addition, DD can act as a signaling molecule, enabling cell-cell communication within a diatom population. However, lethal doses initiate a signaling pathway which includes Ca^2+^ transients, nitric oxide production and mitochondria-specific oxidation of the glutathione pool, leading to activation of PCD^12,14,15^. Recently, another diatom-derived infochemical was identified as sterol sulfate, which is produced in the declining growth phase and induces PCD in *Skeletonema marinoi*16. Several studies suggested a possible role for PCD in diatom response to diverse environmental stress conditions, and its important role during bloom demise^13,14,17–21^.

While the signaling, initiation and execution of PCD in metazoans are well described, the PCD biochemical cascade in protists, including diatoms, is underexplored and the molecular machinery involved in this process is yet to be identified. In metazoans, PCD is coordinated and executed by caspases, a family of cysteine-dependent aspartate-directed proteases^22^. While plants and protists lack caspases, they express distant homologues that share the active cysteine-histidine dyad, known as metacaspases (MCs)^23^. In contrast to caspases, MCs act in monomers and cleave their targets after arginine or lysine^24,25^. MCs are divided into four subgroups defined by the arrangement of the short p10 domain, and the catalytic p20 domain (Fig. 1a). A bioinformatics analysis in phytoplankton genomes identified type III MCs, the only type in which the p10 domain precedes the p20 (Fig. 1a). Type III MCs are absent in fungi, plants and green algal lineages, but prevalent in haptophyta, cryptophyta and heterokontophyta (including diatoms)^26^. To date, the role and function of type III MCs in the cell is unknown^27^. Expression levels of MCs in diatoms were correlated with iron and silica limitations that led to the induction of PCD^17,28,29^, but direct functional or biochemical evidence regarding the role of diatom MCs in PCD and acclimation to stress are yet to be described.

**Fig. 1.**
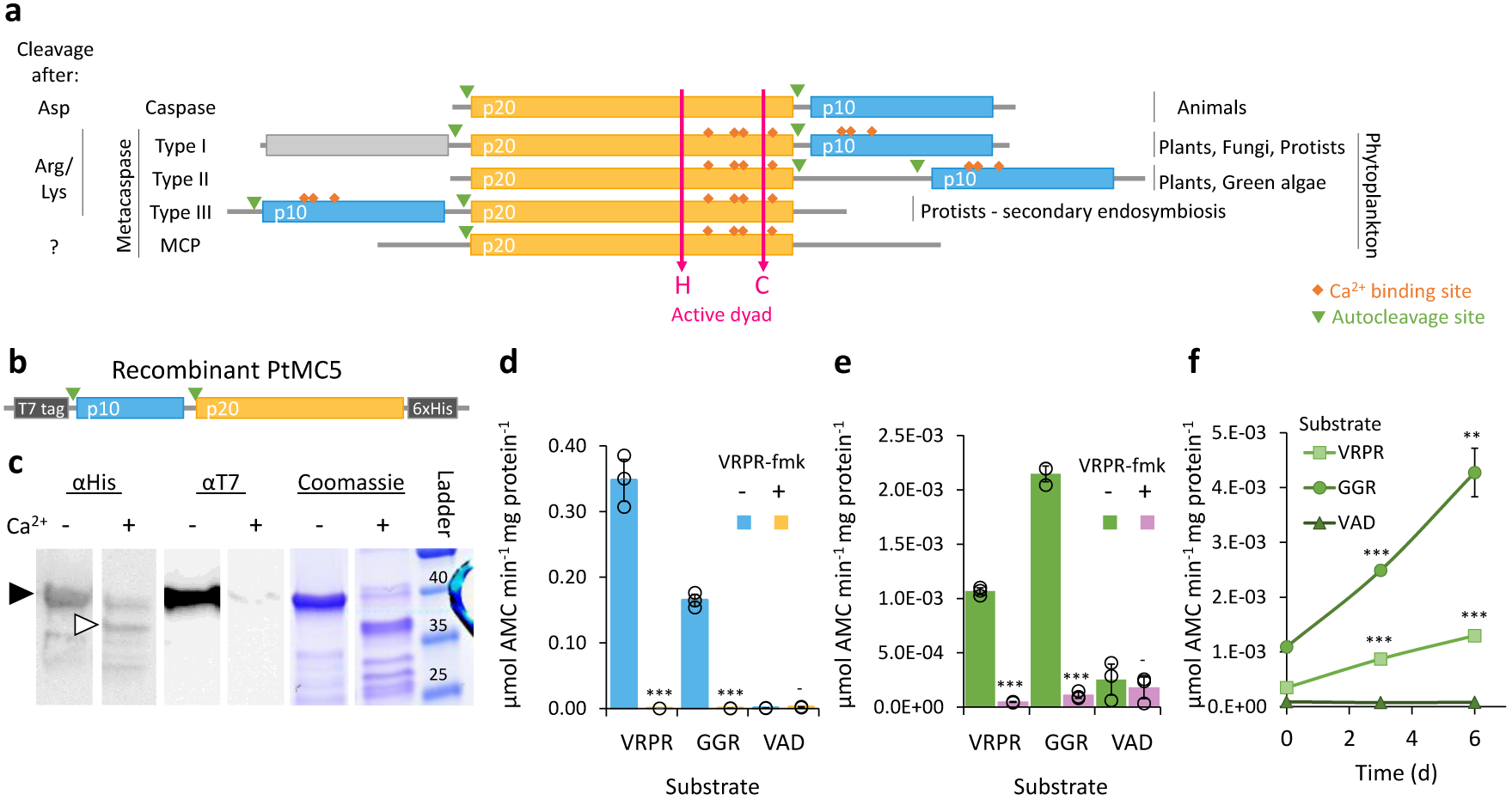
PtMC5 exhibits Ca^2+^ dependent autocleavage and MC-typical protease activity. **a** Domain organization of caspases, MCs, and MC-like proteases (MCPs), as previously described26,36. The p20 and p10 domains are marked in orange and blue respectively. Autocleavage sites are marked with green triangles, Ca^2+^ binding sites in orange rhombuses, the active His-Cys dyad in magenta arrows. **b** Schematic representation of the recombinant tagged PtMC5 expressed in *E.coli*. **c** Coomassie stained SDS-PAGE and immunoblot with antibodies against His or T7 tags. About 2.5 μg purified protein extracts per lane were incubated with (+) or without (-) 10 mM CaCl_2_ for 30 min prior to gel loading. Black arrowhead represents the full length PtMC5, white arrowhead represents auto-cleaved PtMC5 as detected following Ca^2+^ addition. Protein ladder (size in kD) is presented in the right. **d**, **e** Protease activity of purified recombinant PtMC5 (**d**) and of protein extracts from exponential *P. tricornutum* cells (**e**), measured by the release of AMC from peptidyl substrates, with (+) or without (-) 25 μM of the MC inhibitor VRPR-fmk. Standard curve was used to convert the relative fluorescence units into μmol of free AMC released per min per mg of total protein. **f** Protease activity of protein extracts from *P. tricornutum* cells at different time points along the growth curve, measured as in **e**. Single measurements are indicated in circles, bars are means ± s.d. of triplicates, compared to no inhibitor (**d**, **e**) or to earlier time point (**f**)-*P* >0.05,^**^*P*<0.005, ^***^*P* <0.001.

In this study, we combined biochemical characterization of a recombinant type III MC from the model diatom *Phaeodactylum tricornutum* (PtMC5), with functional characterization of genetically modified *P. tricornutum* cells, in order to unveil the function and role of MCs in diatom cell fate regulation. We demonstrate that *PtMC5* encodes an active Ca^2+^ dependent Cys-protease, and identified a unique redox regulation of MC activity by oxidation of two regulatory Cys. This regulatory Cys couple is specific to diatom type III MCs, in which it is highly prevalent. We further demonstrate that PtMC5 mediates cell death in response to DD, suggesting the role of 2-Cys MCs in diatom PCD.

## Results

### *In vitro* characterization of PtMC5 biochemical activity

The *P. tricornutum* genome encodes for 5 MCs: PtMC1 and PtMC3 are MC-like protease (MCPs, see Fig. 1a), their relative expression levels were very low under various conditions (data obtained from published transcriptomes^30,31^, Supplementary Fig. 1), and the proteins were not detected (data obtained from published proteomes^32–34^). In contrast, the type III MCs, PtMC2, PtMC4, and PtMC5 proteins were detected^32–34^, and had higher gene expression levels in all the examined conditions. The gene expression levels in steady state, and the protein abundance of all the type III MCs were similar. We chose to focus on PtMC5, which was highly expressed in 3 independent transcriptomes under diverse growth phases, and induced in stress conditions, including transition to the dark, nitrogen limitation and phosphate limitation^30,31,35^ (Supplementary Fig. 1). Using protein sequence alignment, we detected conserved Ca^2+^ binding sites in PtMC5, both in the p20 and in the p10 domains, and putative auto-cleavage sites between the p10 and p20 domains and before the p10 domain (Supplementary Fig. 2). To characterize its biochemical function, we heterologously expressed a recombinant T7 and 6xHis tagged PtMC5 in *Escherichia coli* cells (Fig. 1b). A full-length PtMC5 protein was detected by a distinct ~40 kD band on SDS-PAGE and immunoblots using antibodies either against the C-terminus T7 tag or the N-terminus 6xHis tag, confirmed by the expression of a full-length PtMC5 protein (40 kD) in *E. coli* (Fig. 1c, black arrowhead). Incubation of PtMC5 with 10 mM CaCl_2_ for 30 min led to autoprocessing and revealed several new bands. A ~37 kD band (Fig. 1c, white arrowhead) may represent the cleavage before the p10 domain (positions 5-6), whilst cleavage between the p10 and p20 domains (positions 122-123) may lead to the ~25 kD band (Fig. 1c, Supplementary Figs. 2 and 3).

*In vitro* activation of MCs was shown to require millimolar concentrations of Ca^2+^, which binds to the Ca^2+^ binding site, and dithiothreitol (DTT), a reductant essential for the reactivity of the active-site Cys^37,38^. Under reduced conditions, PtMC5 exhibited a MC-typical activity, i.e. calcium-dependent cleavage after arginine, monitored as GGRase activity (cleavage after the short peptide Gly-Gly-Arg). This reached saturation at 10 mM Ca^2+^ and was dependent on DTT concentration (Supplementary Fig. 4). PtMC5 displayed preferential VRPRase activity, which was 2-fold higher than its GGRase activity. In contrast, PtMC5 exhibited no caspase-typical activity, as cleavage of the pan-caspase substrate z-VAD-AMC was ~3 orders of magnitude lower than cleavage of VRPR-AMC (0.0007±0.0002 and 0.3475±0.0320 μmol AMC·min^-1^·mg protein^-1^ respectively, Fig. 1d). PtMC5 activity was completely abolished by the MC inhibitor z-VRPR-fmk (25 μM), and was unaffected by the pan-caspase inhibitor z-VAD-fmk (100 μM, *P* =0.43) (Fig. 1d, Supplementary Fig. 5a). Together, these results demonstrate that PtMC5 exhibits a Ca^2+^ dependent MC-typical activity and does not exhibit caspase-typical activity.

Following *in vitro* biochemical characterization, we examined whether PtMC5 typical activity could be also detected in cell extracts of *P. tricornutum*. Similar to the recombinant PtMC5, *P. tricornutum* cell extracts exhibited typical MC activity, showing cleavage after arginine, albeit with preference to GGRase, over VRPRase activity. This MC-typical activity was an order of magnitude higher than caspase-typical activity (VADase) (Fig. 1e). In accordance, MC-typical activity, but not the VADase activity, was inhibited by the MC inhibitor z-VRPR-fmk (25 μM, Fig. 1e, pink bars). The caspase inhibitor z-VAD-fmk (100 μM), inhibited VRPRase activity by ~20% (*P* =0.004), but did not affect the VADase activity (*P*=0.430) (Supplementary Fig. 5b). This demonstrates MC-typical activity in *P. tricornutum* cell extracts, which is likely derived from the combined activity of PtMCs and additional proteases.

Since PtMC5 transcription was shown to be induced during culture growth (data from Matthijs *et al.*^30^ Supplementary Fig. 1), we monitored MC-typical activity during 8 days of growth. On day 3 (exponential phase), GGRase and VRPRase activity exhibited a 2.3 and 2.5 fold increase respectively, compared to day 0 (*P*=0.00014 for both, Fig. 1f). MC-typical activity reached a maximum in early stationary phase, with a 1.7 and 1.5 fold increase in day 6 compared to day 3 of GGRase (*P*=0.0050) and VRPRase (*P*=0.0049) activity respectively.

### Functional characterization of PtMC5 in *P. tricornutum* cells

In order to assess the functional role of PtMC5 in regulating *P. tricornutum* cell fate, we either overexpressed PtMC5, or deleted the active site using CRISPR/Cas9 (Supplementary Fig. 6a-c). Two independent transformant lines for overexpression (OE7, OE9) and knockout (KO1, KO3) were selected after verification using PCR screening of the *PtMC5* gene. The overexpressed *PtMC5* had a shorter band as expected (no introns) in addition to the endogenic *PtMC5* in the OE lines (Fig. 2a). High expression of *PtMC5* in OE lines was verified by real-time qPCR (Fig. 2b). In the KO lines, an edited PtMC5 was detected, indicating a bi-allelic deletion of ~100 bp (Fig. 2a). Exact deletions were assessed by DNA sequencing (Supplementary Fig. 6d). Importantly, in the two KO lines, PtMC5 lacked the putative catalytic Cys (C264), and the deletion led to a frame-shift and an early stop codon (Supplementary Fig. 6e). VRPRase activity in the *P. tricornutum* cell extracts of the transformant lines was 2.2-3.3 fold higher in OE lines compared to WT, and 0.3-0.7 fold lower in KO lines compared to WT (Fig. 2c), indicating that PtMC5 is responsible for at least part of the VRPRase activity detected in cell extracts. Nevertheless, the growth rate of all the transformant lines was comparable to WT, although KO lines reached slightly lower cell concentrations in stationary phase (at day 7 KO1, *P*=0.0001, KO3, *P*=0.0055, Supplementary Fig. 7a).

**Fig. 2.**
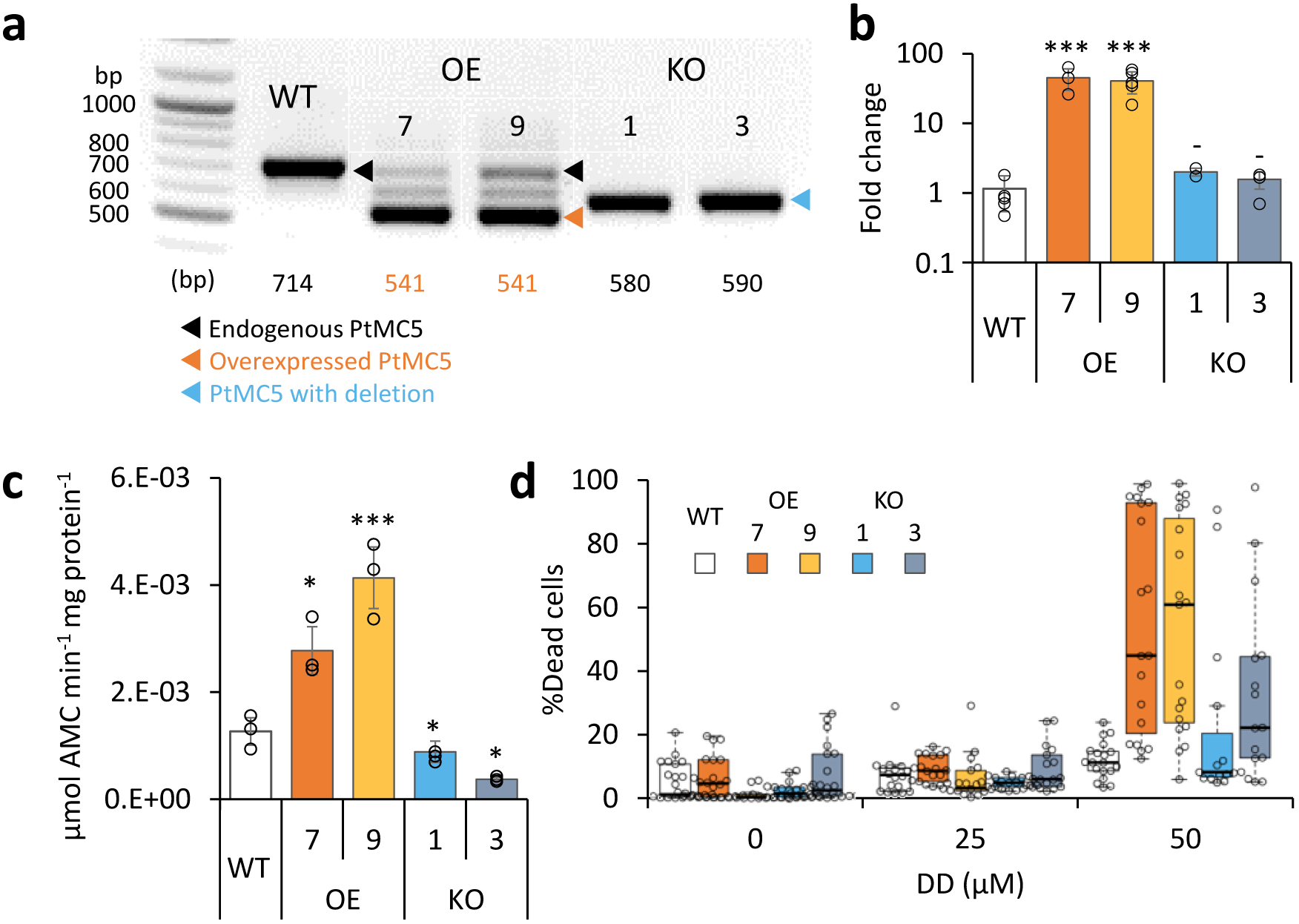
PtMC5 overexpression and knockout in *P. tricornutum* cells. **a** PCR of the *PtMC5* gene in WT, overexpression (OE) and knockout (KO) lines. DNA ladder (size in bp) is present in the left lane, the predicted band size is indicated below. OE lines were generated using a cDNA construct without the introns, hence the shorter band, in addition to the endogenous *PtMC5*. Homozygous deletion events, induced by CRISPR/Cas9 directly evidenced by the presence of a single, shorter PCR product (KO1, KO3), compared to WT cells. **b** Expression levels of PtMC5 normalized to TATA box Binding Protein (TBP), and to WT cells, measured by RT-qPCR, in WT, OE and KO lines. Single measurements are indicated in circles, bars are means ± s.d of biological triplicates. **c** PtMC5 VRPRase activity in protein extracts of WT, OE, and KO *P. tricornutum* lines. Single measurements are indicated in circles, bars are means ± s.d. of triplicates. **d** Box plot of % Sytox positive cells of cultures treated with 0, 25, 50 μM DD at early stationary phase. Each point represents a single replica. Data was obtained from 6 independent experiments, each including 3-4 biological replicates. Colored boxes – 1st to 3rd quartiles; horizontal line within the box – median, N = 19. Flow cytometry analysis was based on fluorescent measurements of at least 10,000 cells per sample. Each transformant line was compared to WT-*P*>0.05, ^*^*P*<0.05,^**^*P*<0.01, ^***^*P*<0.005.

To examine whether PtMC5 has a role in cell fate regulation, we monitored the survival of WT and PtMC5 transformant lines following treatment with the diatom-derived infochemical DD, which was shown to induce PCD hallmarks in *P. tricornutum* in a dose-dependent manner^12,13,15,39^. Since MC-typical activity was induced in early stationary phase, during which KO lines exhibited lower cell abundance, we examined the responses of the different strains to DD in early stationary phase (day 6). In a collection of 6 independent experiments of early stationary cultures, treatment of WT cells with 50 μM DD led to 11.2% (median) dead cells, while both OE lines displayed a significantly higher fraction of dead cells (Fig. 2d, median 45.0% and 61.1%, *P*=0.012 and 0.002 in OE lines 7 and 9 respectively). The same trend of higher morality in OE lines compared to WT was repeated in 8 experiments (Supplementary Fig. 7b). The response of the KO lines was inconsistent, in some experiments they were more resistant than WT cells, whilst in others they were more sensitive, exhibiting high variance within biological triplicates. No significant difference in cell death was detected in exponential cultures of WT and transformant lines after DD treatment (Supplementary Fig. 7c). Taken together, these results point towards an important role for PtMC5 activity as a positive regulator of DD-induced PCD in *P. tricornutum*, specifically during stationary phase.

### C264 is essential for PtMC5 catalytic activity, while C202 is a regulatory Cys

The activity of proteins involved in executing PCD ought to be tightly regulated, especially when the proteins are basally expressed as PtMC5; therefore PCD executers are often present as inactive zymogens. Protein activity is often regulated by reversible Cys oxidation, where the oxidation can induce or inhibit the enzymatic activity. Based on our previous work exposing the redox proteome of *P. tricornutum* (redoxome)^33^, we could detect redox-sensitive cysteines in PtMCs that showed significant oxidation upon treatment with lethal doses of DD (25 μM) and H_2_O_2_ (150 μM), in comparison to non-lethal treatments (5 μM DD 2 h prior to 25 μM DD, and no H_2_O_2_)^33^. Degree of cysteine oxidation was measured for each detected peptide, and delta oxidation was calculated by subtraction of the oxidation degree of the non-lethal treatments from the lethal treatments. *P. tricornutum* MCPs (PtMC1, PtMC3) were not detected, in accordance with their low RNA expression levels (Supplementary Table 1), while peptides representing all type III MCs were detected (Table 1, Supplementary Table 1). Cys 144 in PtMC5 and its homologues in PtMC2 and PtMC4 were detected, but did not undergo significant oxidation due to lethal treatments (Table 1, Supplementary Table 1, and Fig. 3a, red frames). An additional cysteine in PtMC5, C202, and its homologues in PtMC4 were detected in the two redox proteomes. In PtMC5 the oxidation was significantly higher in response to H_2_O_2_ and even higher in response to DD, exhibiting 20.9% more oxidation in the lethal DD treatment compared to the non-lethal treatment (Table 1, Supplementary Table 1). Out of 5 detected cysteines in 3 MCs, only C202 was significantly oxidized in response to lethal treatments, suggesting that its oxidation is specific, and has a possible involvement in regulating PtMC5 activity.

**Table 1.**
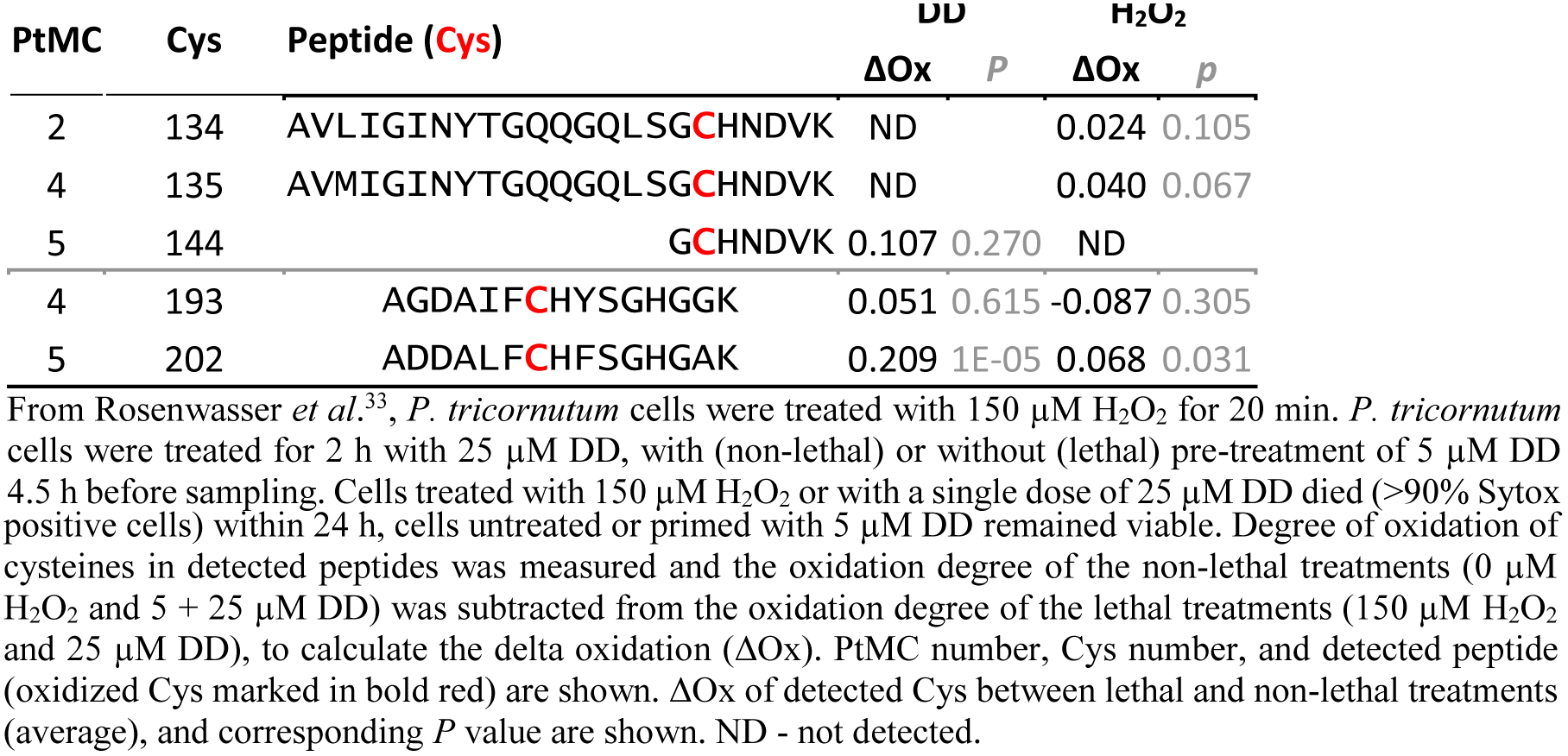
All PtMCs peptides detected by redox proteomics in response to DD or H_2_O_2_ treatments.

**Fig. 3.**
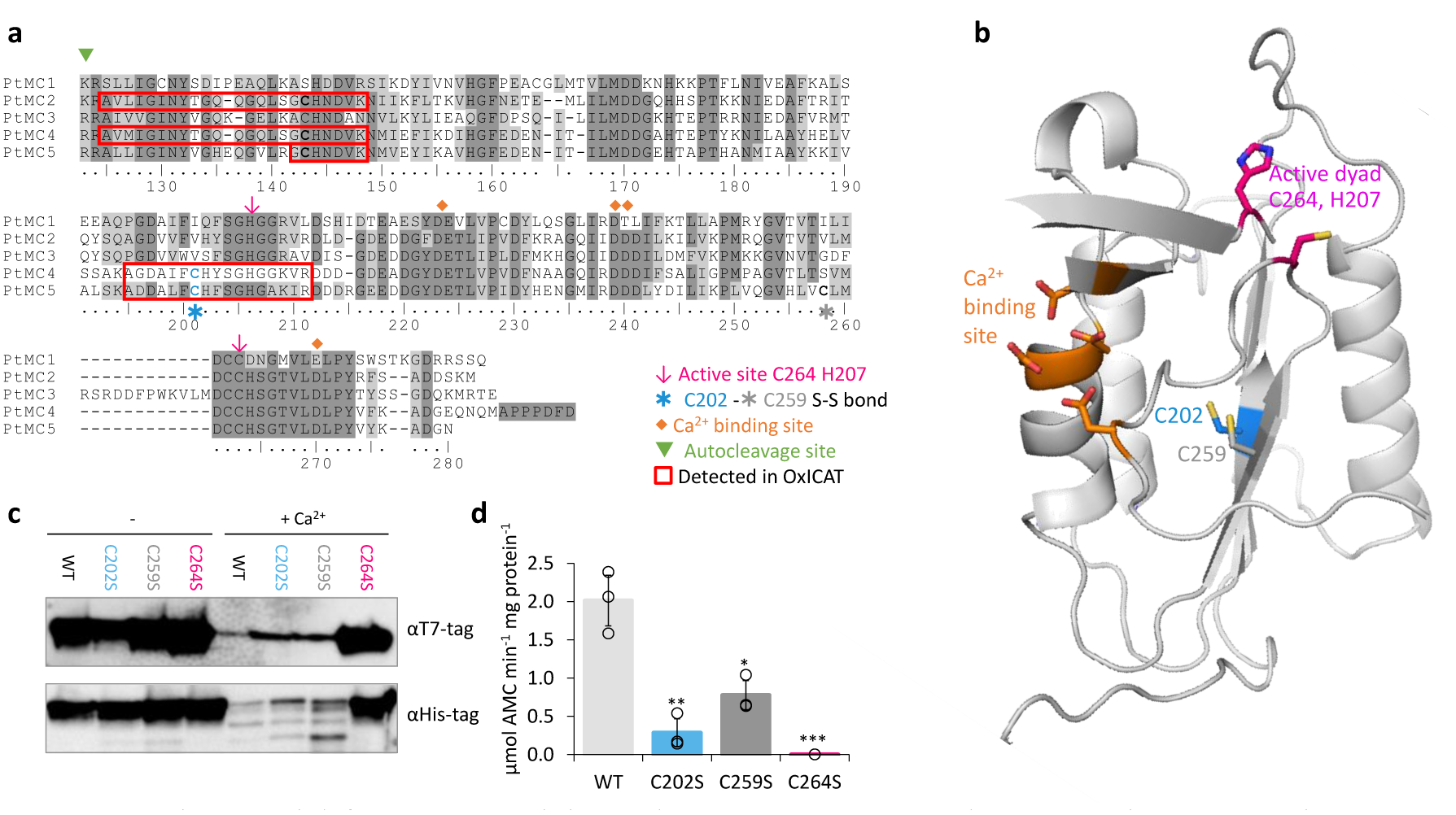
C264 is essential for PtMC5 activity and C202, C259 are regulatory cysteines. **a** Protein sequence alignment of PtMCs p20 domains, amino-acids numbered by PtMC5 sequence. Identical residues have dark gray background and similar amino acids have light gray background 70% threshold for coloring. Active dyad, C202, C259, Ca^2+^ binding site, and autocleavage sites are marked. Peptides detected in redox proteomics are framed in red. **b** PtMC5 3-dimentional structure model based on *S. cerevisiae* MC structure^41^, cartoon representation by PyMOL. Key amino acids are presented in stick view, Sulfur, Oxygen and Nitrogen atoms marked conventionally in yellow, blue and red respectively. Aspartates of the Ca^2+^ binding site are marked in orange, the active Cys histidine dyad is marked in magenta, C202 which was detected in redox proteomics is marked in blue, neighboring C259 is marked in dark gray. **c** Immunoblot with (HRP) αHis or αT7 tags of PtMC5, PtMC5^C202S^, PtMC5^C259S^, and PtMC5^C264S^. About 0.88 μg protein extracts per lane were incubated in activity buffer without / with 10 mM CaCl_2_ for 10 min prior to gel loading. **d** Recombinant PtMC5, PtMC5^C202S^, PtMC5^C259S^, and PtMC5^C264S^ protease activity. Single measurements are indicated in circles, bars are means ± s.d. of triplicates. Mutants lines were compared to WT, ^*^*P*=0.010,^**^*P*=0.003, ^***^*P*=0.001.

Since C202 is not conserved in other organisms (data from Woehle *et al*.^40^), and there is no known redox regulation of MC activity, a structural based analysis was performed *in silico* to examine the potential function of the C202. We used the *Saccharomyces cerevisiae* MCA1 structure^41^ (a type I MC) as a basis for the PtMC5 structural model. The model captured the p20 domain structure (Supplementary Fig. 8) and indicated that C202 is distant from the active site, located in the core of the protein 4.1 Å apart from C259. This distance is within the range of a reversible disulfide bond^42,43^, which upon oxidation may form between the two β-sheets, hence stabilizing the protein (Fig. 3b). Notably, this cysteine pair is unique to PtMC5, and is absent in PtMC1-4 (Fig. 3a), and the *S. cerevisiae* MC (Supplementary Fig. 8).

To investigate the function of C202, mutant forms PtMC5^C202S^ and PtMC5^C259S^, in which the Cys was substituted with Ser, thus eliminating the potential formation of the disulfide bond, were overexpressed in *E. coli* and tested for their activity. In addition, we tested PtMC5^C264S^, in which the putative catalytic Cys was mutated. The mutant in the active-site, PtMC5^C264S^ exhibited a loss of activity as expected, with no apparent autocleavage activity and 3 orders of magnitude lower VRPRase activity compared to the WT (Fig. 3c, d). The mutants PtMC5^C202S^ and PtMC5^C259S^, without the putative disulfide bond, were still active and were able to undergo autoprocessing (Fig. 3c). However, the autocleavage rate appears to be slower, as the ~40 kD band was still apparent after 10 min activation but disappeared after 30 min activation (Fig. 3c, Supplementary Fig. 9). Furthermore, PtMC5^C202S^ and PtMC5^C259S^ recombinant proteins exhibited lower VRPRase activity compared to the WT (15% and 40%, *P*=0.003, *P*=0.010 respectively, Fig. 3d). Thus, this data supports a regulatory role for the suggested disulfide bond between C202 and C259 (herein 2-Cys), while C264 is essential for PtMC5 proteolytic activity.

### The 2-Cys type III MCs are prevalent and specific to diatoms

We aligned the p20 domain across diverse photosynthetic organisms and protist species in order to map the abundance of 2-Cys MCs (Table 2). The 2-Cys were absent in green and red algae, glaucophytes, cryptophytes, haptophytes and alveolates. In the group of stramenopiles, only diatoms were found to encode for 2-Cys MCs. Importantly, several diatom species with global distribution, such as the centric bloom-forming *S. marinoi* and *Thalassiosira pseudonana* possess 2-Cys MCs (Table 2).

**Table 2.**
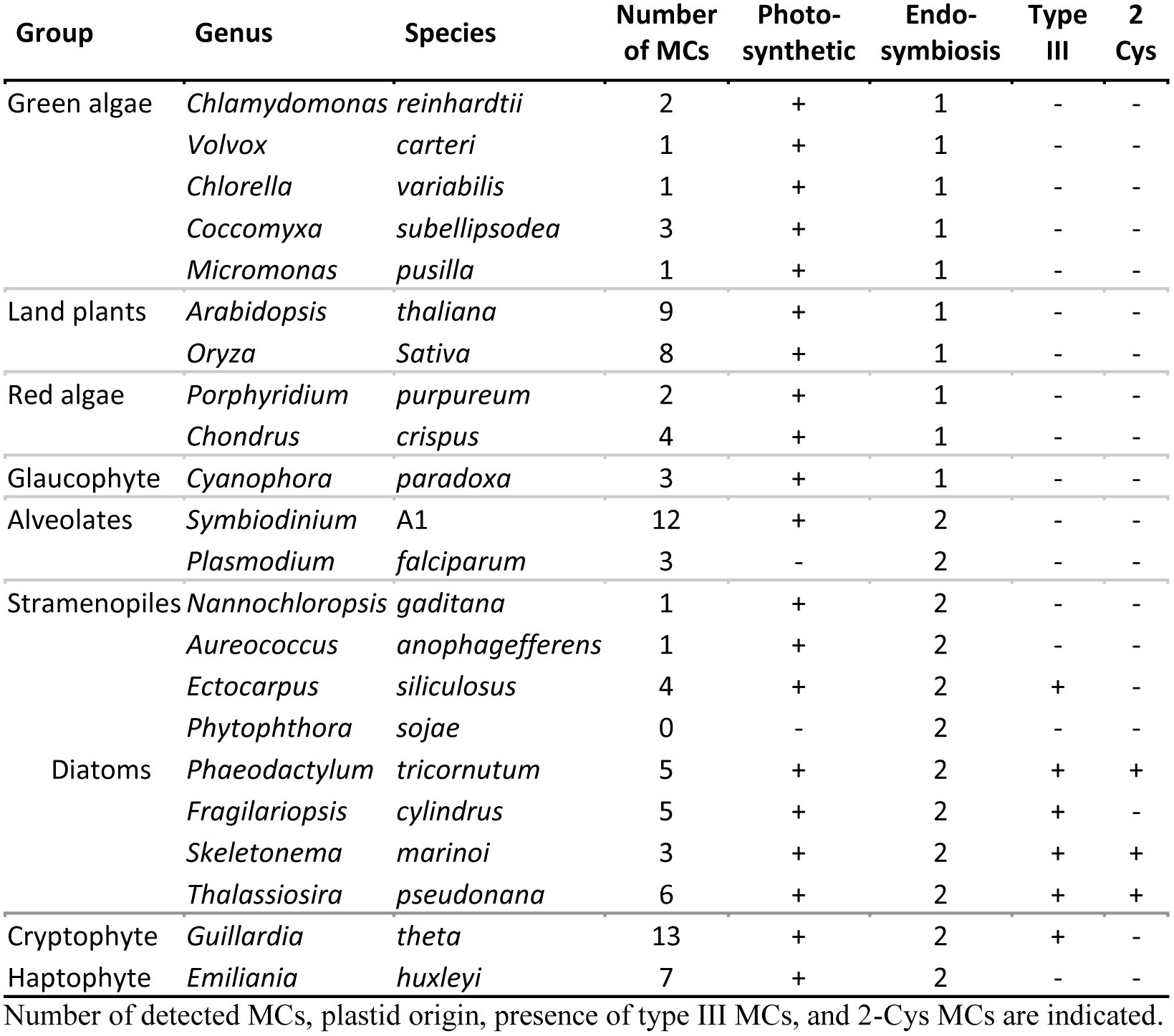
Abundance of 2-Cys MCs across species.

Furthermore, we identified 2-Cys MCs in an additional 19 diatom species (Supplementary Table 2) based on the Marine Microbial Eukaryote Transcriptome Sequencing Project (MMETSP)^44^. The high relative abundance (62%) of 2-Cys MCs in diatoms of the MMETSP dataset indicates that 2-Cys MCs are not only widely distributed in diatoms but also widely expressed. Remarkably, all the 2-Cys are type III MCs. So far all type III MCs have only been found in secondary endosymbionts^26^. Hence, we defined 2-Cys MCs as a novel subtype of type III MCs. Phylogenetic analysis of the 134 diatom MCs, based on the p20 domain, revealed that the 2-Cys were probably acquired in a few independent events (Fig. 4, Supplementary Fig. 10). PtMC5 belongs to a clade of 4 MCs from pennate diatoms, another 5 MCs harboring the 2-Cys are clustered with different clades. The majority (19) of the 2-Cys MCs are from polar-centric diatoms, and clustered together, suggesting a common origin.

**Fig. 4.**
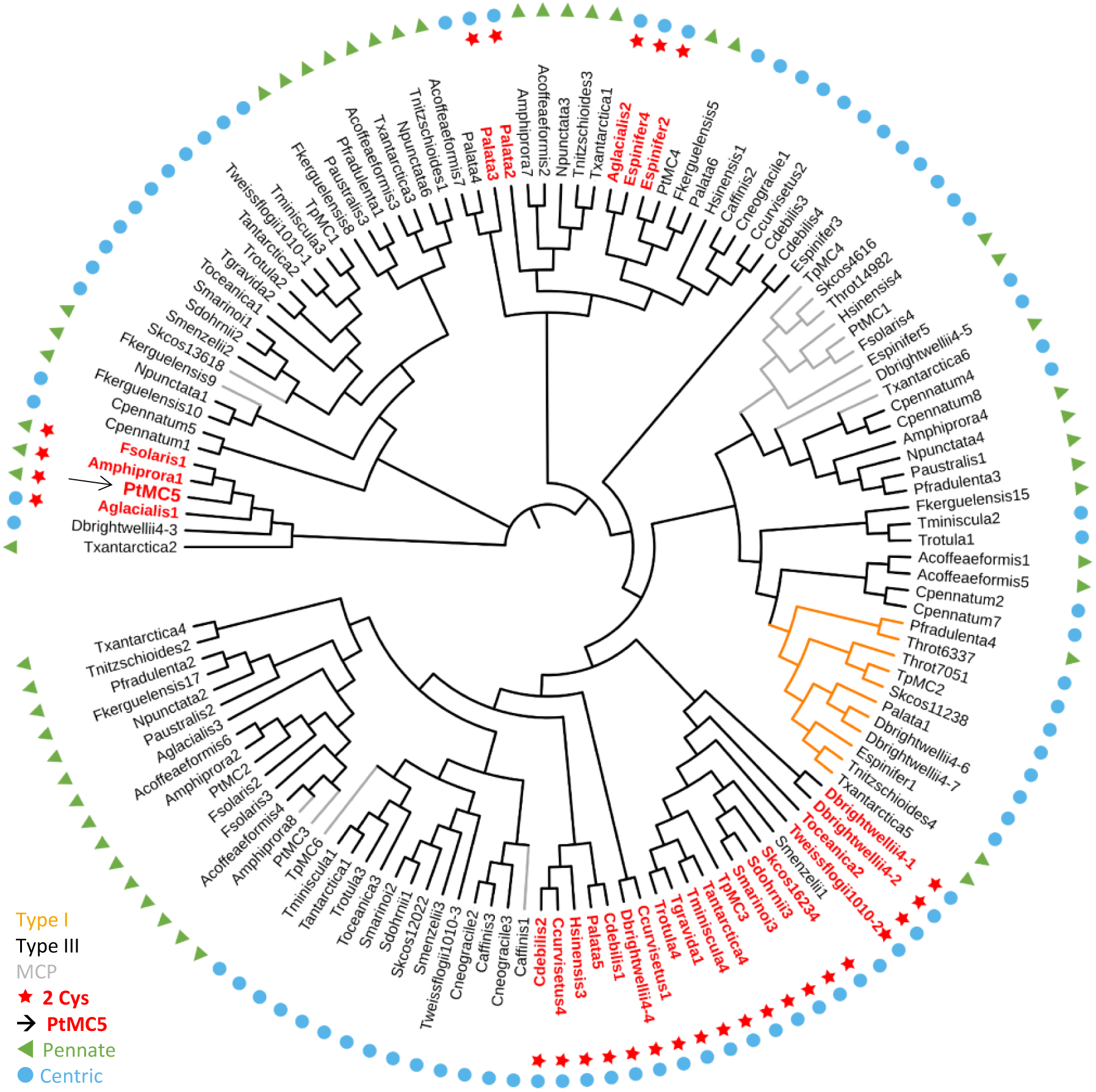
A phylogenetic tree of diatom MCs based on protein sequence alignment of the p20 domain identified in the MMETSP dataset^44^. Type I MCs are marked in orange lines, MCPs are marked in gray lines, and type III MCs are marked in black lines. 2-Cys MCs are marked with red bold text and stars, PtMC5 is marked with a black arrow. Pennate diatoms are marked with green triangles, centric diatoms with blue circles. Numbers indicate the MC number, *P. tricornutum* and *T. pseudonana* MCs are numbered according to Choi^26^.

Based on metatranscriptome analysis of a natural diatom bloom dominated by *Skeletonema* species and *T. rotula* in Narragansett Bay^45^, we further confirmed expression of 2-Cys type III MCs in a natural diatoms community (gene IDs 16234 and 16314 respectively, Supplementary Fig. 11). To summarize, the newly defined 2-Cys MC subtype of type III MCs appears to be diatom specific and is expressed in laboratory experiments and a natural diatom blooms.

## Discussion

In the last two decades numerous studies have reported hallmarks of PCD that are prevalent in a wide range of microorganisms including bacteria, yeast, protozoans and diverse phytoplankton groups^10,21,46^. Though the molecular pathways of PCD have not yet been characterized in phytoplankton, MCs were suggested as possible PCD regulators, due to their structural similarity to the canonical caspases, which are the metazoan PCD regulators and executers. Furthermore, MCs were also shown to participate in PCD activation in plants in various tissues and at different developmental stages^47^ as well as in response to stress^48^. To date, none of the putative PCD proteins in phytoplankton, including MCs, have been functionally characterized, although several studies have monitored MCs expression under different environmental stress conditions which can lead to PCD^17,20,29,49,50^.

In this study, we characterized for the first time the biochemical function, regulation and ecophysiological significance of a diatom type III MC, PtMC5, belonging to a novel subtype of 2-Cys type III MCs that appears to be unique to diatoms. Recombinant PtMC5 exhibited MC-typical activity - calcium dependent autoprocessing and cleavage after arginine. Although caspase-typical activity (cleavage after aspartate) was previously detected in diatoms^17,28,29,51^, we show that PtMC5 does not exhibit caspase-typical activity, and in *P. tricornutum* the caspase-typical activity is an order of magnitude lower than the MC-typical activity. In contrast to recombinant PtMC5, *P. tricornutum* cells exhibited higher GGRase than VRPRase activity (Fig 1d, e), probably due to combined activity of PtMC5 with additional PtMCs and other proteases. Accordingly VRPRase activity, representative of PtMC5 activity was enhanced in *P. tricornutum* cells overexpressing PtMC5 and decreased in PtMC5 knockout lines (Fig. 2c), indicating that part of the MC typical activity detected in *P. tricornutum* cell extracts is indeed the result of the *PtMC5* gene product. We also found that MC typical activity (GGRase as well as VRPRase) was induced during early stationary phase, suggesting that MCs role is specific to the a cell physiological state. In addition, PtMC5 KO lines reached slightly, but significantly lower cell abundance at early stationary phase compared to WT or OE lines (Supplementary Fig. 7a), suggesting a vital role for PtMC5 in growth phase transition. Together with the induction in PtMC5 gene expression^52^ and general MC-typical activity (Fig. 1f), these findings implicate an important role of PtMC5 activity in early stationary phase. Intriguingly, in early stationary phase OE PtMC5 lines exhibited enhanced DD-induced cell death (Fig. 2d, Supplementary Fig. 7b), while in exponential phase, in which PtMC5 expression and MC-typical activity were lower, the survival of both OE and KO lines was similar to WT cells. The lack of a clear phenotype in the KO lines may be a result of redundancy by the other PtMCs which can compensate PtMC5. Furthermore, PCD in unicellular organisms is expected to be a highly regulated process, executed only under very specific stress conditions in subpopulations^53^.

PCD related proteins require very tight regulation on their activation, and execute cell death only upon requirement. In multicellular organisms, the PCD executing caspases are translated as inactive zymogens, and are activated only by a complex biochemical activation cascade, including dimerization and cleavage. The plants and protist homologues, MCs, are activated by Ca^2+^ binding and autoprocessing, and are active as monomers^54^. In accordance, recombinant PtMC5 exhibited Ca^2+^ dependent MC typical activity with a conserved Ca^2+^ binding site in the p20 domain, similar to GtMC2, a type III MC from the cryptophyte *Guillardia theta*^55^. The role of PtMC5 in DD induced PCD is further supported by *in vivo* measurements of a Ca^2+^ burst in response to lethal DD treatment in *P. tricornutum* cells^12^. PtMC5, as seen in other MCs^56,57^, may undergo autolysis (low molecular weight bands Supplementary Figs. 3, 9) as a self-inactivation mechanism, that ensures that the activated PtMC5 will have a short functional half-life. Importantly, we identified another layer of posttranslational regulation, novel in MCs, through a reversible oxidation of reactive regulatory cysteines. By combining data from redox proteomics with a 3D protein model and directed point mutations, we suggest that oxidation of C202, as detected in response to lethal treatments (Table 1), forms a stabilizing disulfide bond with C259, enhancing PtMC5 activity. Mutation in either one of the 2-Cys decreased PtMC5 activity to 15-40% of WT activity (Fig. 3d). In contrast, oxidation of the active site cysteine inactivates plant MCs^37^. Under such a scenario, only mild oxidative stress caused by environmental conditions could lead to specific oxidation of these regulatory Cys, enhancing PtMC5 activity. This tight regulation of protein activity at the post-translational level can allow basal expression and quick activation without *de-novo* protein synthesis.

We propose a model (Fig. 5) in which optimal activation of PtMC5 requires the combination of two signals, Ca^2+^ and a mild reactive oxygen species (ROS). The ROS signal can be generated as an enzymatic byproduct of Ca^2+^ signaling^58^ or as a direct result of an environmental stress. Diverse environmental stresses, including diatom-derived infochemicals such as DD, are perceived by the induction of Ca^2+^ intracellular transients within seconds^12,59^. Subsequently, Ca^2+^ can bind to the MCs Ca^2+^ binding site, which is necessary for MC activation^55,60^ (Fig. 1b, Supplemental Fig. 4a). In addition, DD leads to Ca^2+^ transients followed by a ROS burst in the mitochondria^14,15^. This ROS accumulation is essential to induce the PCD cascade, since addition of antioxidants can prevent the subsequent activation of PCD^15,53,61^. Sublethal ROS levels can act as a signal and oxidize the 2-Cys to form a disulfide bond, which induces PtMC5 activity (Fig. 5). These rapid post-translational modifications lead to activation of pre-existing PtMC5 protein, and execution of a PCD pathway. Only integration of the two signals, Ca^2+^ and ROS, leads to sufficient activity of 2-Cys MCs, decreasing the chance of accidental activation, which may cause unnecessary cell death.

**Fig. 5.**
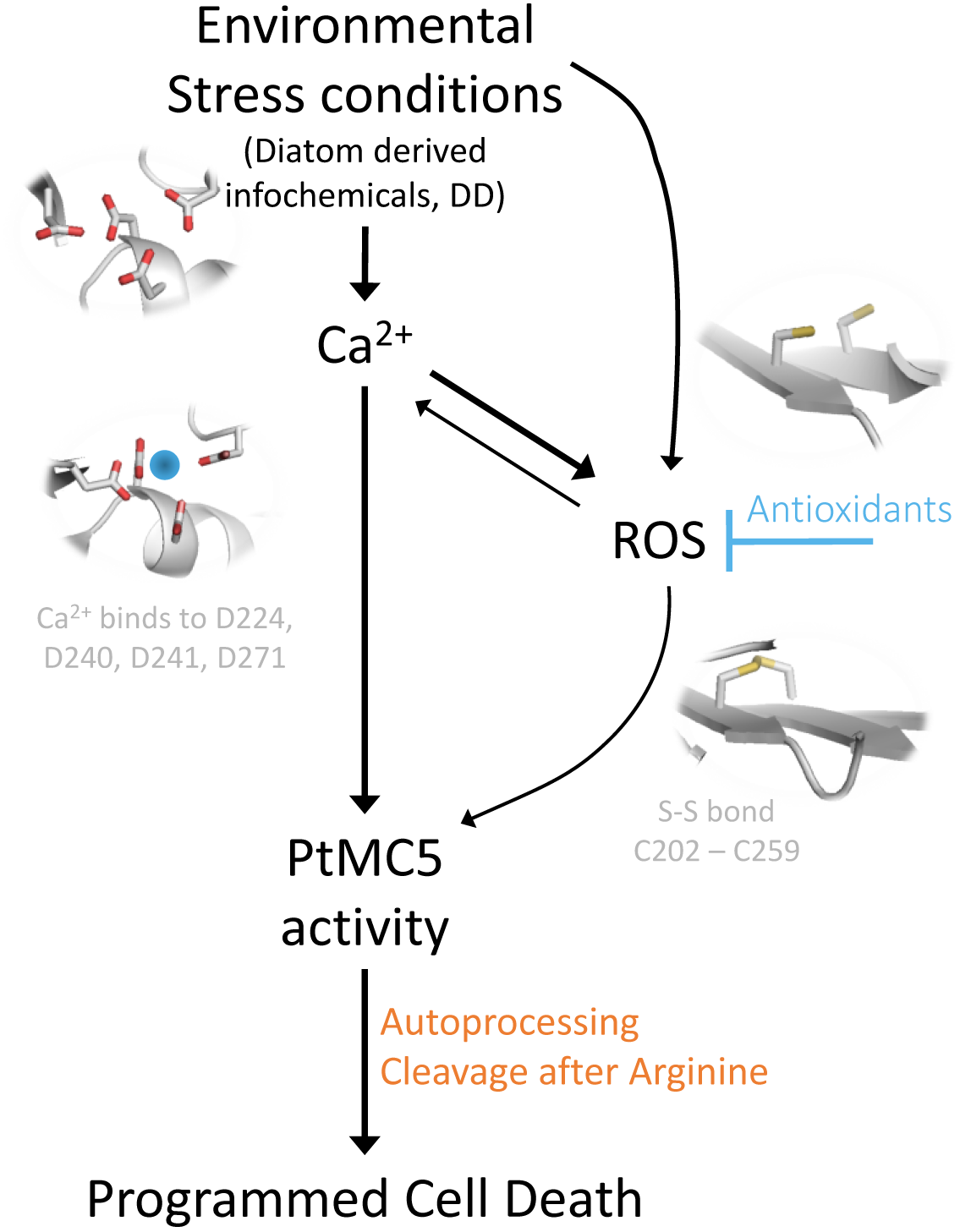
Conceptual model for diatoms’ perception of stress and PCD activation. Environmental stress conditions, such as DD, induce elevation in intercellular Ca^2+^, which leads to oxidative stress and accumulation of ROS. In addition, environmental stresses may cause ROS accumulation directly, which in turn may also lead to Ca transients. The Ca^2+^ binds to the MC Ca^2+^ binding site (Asp 224, 240, 241, 271), and activates the MC. The oxidative stress leads to the formation of a disulfide bond between Cys 202 and 259, enhancing MC activity. PtMC5 activity of autoprocessing and cleavage of target proteins after arginine leads to PCD. The 2-Cys and the Ca^2+^ binding sites were drawn based on the PtMC5 3D model. Oxygen and Sulfur atoms are marked in red and yellow respectively. The calcium atom is marked by blue circle.

Recent phylogenomic analysis tracked the evolutionary history of the redox-sensitive Cys residues in *P. tricornutum*, revealing its expansion during plastid evolution^40^. Interestingly, the unique presence of the 2-Cys in diatom MCs, but not in closely related groups (Supplementary Table 2), suggests a late Cys gain in evolution. Importantly, 2-Cys MCs were expressed in cultures as well as in a natural diatoms bloom (Supplementary Fig. 11). Using a redox-sensitive GFP (roGFP) probe, it was demonstrated *in vivo* that early oxidation of the *P. tricornutum* mitochondrial glutathione pool in response to DD led to PCD in a dose-dependent manner^15^. Redox-regulation on MCs activity via reactive Cys can allow diatoms to integrate various environmental signals in order to rapidly adjust cellular processes in a reversible manner and may also be involved in the activation of the PCD cascade.

In phytoplankton, PCD is correlated to MCs activity and is triggered by diverse environmental stresses, possibly leading to high rates of cell lysis during bloom demise^18,62–65^. Cell lysis in the form of PCD during bloom demise may have an important ecological role in the efficient recycling of essential nutrients and in generating unique metabolic composition that will drive specific microbial interactions^18, 65–67^. Diatoms can form dense, nearly clonal blooms^68^ or form biofilms, thus reaching high local densities that enable effective cell-cell interactions. In such colony-like morphology, PCD can lead to elimination of stressed cells from the population while allowing the survival of the fittest cells and enhancing the survival of the species as a whole. Future studies will help to elucidate the ecophysiological importance of PCD-dependent mortality of phytoplankton blooms and turnover of carbon in the ocean.

## Methods

### Culture growth

*P. tricornutum*, accession Pt1 8.6 (CCMP2561 in the Provasoli-Guillard National Center for Culture of Marine Phytoplankton) was purchased from the National Center of Marine Algae and Microbiota (NMA, formerly known as CCMP). Cultures were grown in f/2 media in filtered seawater (FSW) at 18 °C with 16:8 hours light:dark cycles and light intensity of 80 μmol photons·m^-2^·sec^-1^ supplied by cool-white LED lights. Unless specified otherwise, experiments were initiated with exponentially growing cultures at ~5·10^5^ cells·mL^-1^.

### Cell death

Cell death was determined by positive Sytox Green (Invitrogen) staining, used at a final concentration of 1 μM. Samples were incubated in the dark for 30 min prior to measurement.

### Infochemical preparation

(*E,E*)-2,4-decadienal (DD) (95%, Acros Organics) solutions were prepared by diluting the stock in absolute methanol on ice. DD was added to the cells at a dilution of at least 1:200. Control cultures were treated by the addition of methanol to the same dilution as the treatment culture.

### Flow cytometry

Flow cytometry measurements (cell abundance and Sytox staining) were obtained using the Eclipse iCyt flow cytometer (Sony Biotechnology Inc., Champaign, IL, USA), equipped with 488 nm solid state air cooled 25 mW laser with a standard filter set-up. At least 5000 cells were measured in each sample, with at least 3 biological replicates.

### Identification of redox sensitive cysteines

*P. tricornutum* cells were treated with 5 μM DD, and 2.5 h later 25 μM DD was added to treated (non-lethal) and untreated (lethal) cells. After 2 h cells were sampled by centrifugation of 200 ml per sample. Proteins were extracted and cysteine oxidation was assessed as previously described by Rosenwasser *et al.*^33^. To summarize, proteins were extracted by sonication and the pellet was dried under nitrogen flow to avoid cysteine oxidation. Subsequent to extraction, proteins were dissolved in denaturing buffer (50 mM Tris, pH=8.5 and 0.1% SDS) and subjected to thiol trapping according to the OxICAT methodology^69^ using the cleavable ICAT reagent kit for protein labeling (AB Sciex, Foster City, CA, USA). Downstream proteomics analysis including peptide liquid chromatography, mass spectrometry and data processing were carried out as previously described by Rosenwasser *et al.*^33^

### Bacterial cloning

cDNA of PtMC5 was ordered from GENWIZ (in pUC57), ligated into the bacterial expression vector pET-21a using EcoRI and XhoI restriction sites. This construct was subsequently used as a template for the preparation of PtMC5 mutants (C202S, C259S or C264S), using site directed mutagenesis (SDM). This was carried out using mutagenesis primers 1-6 (listed in Supplementary Table 3) with either KAPA polymerase PCR followed by DpnI digestion or by using a Q5 SDM kit (New England Biolabs, E00554S). Correct ligation and incorporation of mutations was verified by DNA sequencing using primers 7, 8 (Supplementary Table 3).

### PtMC5 expression and purification

MC expression, purification and activity assays were adapted from Bozhkov *et al*.^70^ and McLuskey *et al*.^71^. *E. coli* Rosetta cells were transformed with the expression plasmids and grown in LB containing ampicillin in a shaker at 37 °C. When O.D. reached 0.6, 1 mM IPTG was added for overnight shacking at 16 °C or 20 °C. Cell pellets collected from 100 ml of bacterial culture were resuspended in 3.5 ml of lysis buffer (150 mM NaCl, 25 mM HEPES, 10% glycerol, 0.2% triton, 1 mg ml^-1^ lysozyme, 1 μl benzonase, 0.5 mM DTT, pH 7.8) and sonicated 10×10 sec on ice. Following centrifugation at 14,000 g for 5 min to remove insoluble debris, the supernatant was applied to Ni-NTA resin (Ni-NTA His•Bind^®^ Resin, Milipore, 70666-3). After washing with base buffer (150 mM NaCl, 25 mM HEPES, 10% glycerol), and with base buffer containing 20 and 30 mM imidazole, the bound proteins were eluted in base buffer, containing 200 mM imidazole. The elution was concentrated and washed with base buffer using an Amicon Centrifugal Filter Units (Millipore) equipped with a 10 kDa exclusion membrane.

Protein concentration was determined using the BCA method and the samples were diluted to the lowest concentration in base buffer. Samples were then incubated with or without 10 mM Ca^2+^. Samples were incubated at 95° C for 5 min and loaded on Tris-Glycine eXtended gels (Criterion TGX Gels Any kD, BioRad) and subjected to protein gel analysis using Coomassie brilliant blue, or blotted onto a poly(vinylidene difluoride) (PVDF) membrane and analyzed using HRP-anti-6xHis or HRP-anti-T7 antibodies (Zotal). The ECL-Prime western blotting detection reagent (GE Healthcare) was used for detection.

### Kinetic assays

Purified PtMC5 or *P. tricornutum* cell lysate (10^8^ cells were harvested, resuspended in 250 μl lysis buffer, sonicated, and centrifuged to remove insoluble debris) was used for kinetic measurements. In each experiment the protein concentration was calculated by the BCA method and all samples were diluted in base buffer. Purified PtMC5 was used at about 60 ng per well and *P. tricornutum* protein extracts were used at about 30 μg per well. Protein extracts were incubated in activity buffer for 30 min prior to addition of the substrate. MC-typical activity, cleavage after arginine/lysine, was assessed using the short peptides Val-Arg-Pro-Arg (VRPR) and Gly-Gly-Arg (GGR) conjugated to the fluorophore 7-Amino-4-methylcoumarin (AMC). Following proteolytic activity the fluorophore was released to the media and its fluorescence was detected over time with 360 nm excitation and 460 nm emission using a plate reader (Infinite 200 pro, Tecan). A calibration curve with 12, 6, 3, 1.5, 0.75, 0 μM AMC and initial slopes were used to calculate the activity (μmol AMC·min^-1^·mg protein^-1^) from the fluorescence measurements as previously described^71^. Assays were performed in activity buffer (base buffer with 0.1% CHAPS, 10 mM DTT, 10 mM CaCl_2_) unless otherwise stated, at 20 °C in black 96 well plates (TAMAR). All assays were performed with 50 μM substrate (z-GGR-AMC, Ac-VRPR-AMC, z-VAD-AMC and AMC, all from Bachem). Protease inhibitors, in which the uncleavable fluoromethylketone (fmk) group is conjugated to the short peptide, z-VRPR-fmk (25 μM, MC inhibitor) and z-VAD-fmk (100 μM, pan-caspase inhibitor) (both from abcam) were incubated for 30 min before the addition of the substrate. Importantly, the measurement of protease activity in cell extracts, in adequate activity buffer, does not represent the actual *in vivo* activity as MCs are often inactive zymogens, but the potential of MC typical activity upon activation signal.

### gRNA design for PtMC5 knockout

In order to inactivate PtMC5 we adapted for *P. tricornutum* the method established by Hopes *et al*. for the diatom *T. pseudonana*^72,73^. Two sgRNAs were designed to cut 115 nucleotides, which includes the catalytic Cys and the 3^rd^ intron. Selection of CRISPR/Cas9 targets and estimating on-target score: Twenty bp targets with an NGG PAM were identified and scored for on-target efficiency using the Broad Institute sgRNA design program (www.broadinstitute.org/rnai/public/analysis-tools/sgrna-design), which utilizes the Doench^74^ on-target scoring algorithm. The sgRNAs that were chosen had no predicted off-targets: The full 20 nt target sequences and their 3′ 12 nt seed sequences were subjected to a nucleotide BLAST search against the *P. tricornutum* genome. Resulting homologous sequences were checked for presence of an adjacent NGG PAM sequence at the 3′ end. The 8 nt sequence outside of the seed sequence was manually checked for complementarity to the target sequence. In order for a site to be considered a potential off-target, the seed sequence had to match, a PAM had to be present at the 3′ end of the sequence and a maximum of three mismatches between the target and sequences from the BLAST search were allowed outside of the seed sequence.

### Plasmid construction using Golden Gate cloning

Golden Gate cloning was carried out previously described^75^, in a design similar to Hopes *at al.*^72^ (Supplementary Fig. 6a). BsaI sites and specific 4 nt overhangs for Level 1 (L1) assembly were added through PCR primers. Golden Gate reactions for L1 and Level 2 (L2) assembly were carried out, 40 fmol of each component was included in a 20 μl reaction with 10 units of BsaI or BpiI and 10 units of T4 DNA ligase in ligation buffer. The reaction was incubated at 37 °C for 5 h, 50 °C for 5 min and 80 °C for 10 min. Five μl of the reaction was transformed into 50 μl of NEB 5α chemically competent *E. coli*.

Level 0 assembly: The endogenous FCP promoter FCP terminator and the Ble resistance gene were amplified from the PH4-pPhat plasmid, and the U6 promoter^76^ from gDNA, using primers 9-14 and 19-20 (Supplementary Table 3). Both promoters are associated with high expression levels. Products were cloned into a pCR8/GW/TOPO vector (ThermoFisher). FCP promoter and terminator were “domesticated” to remove the Bpi sites using a Q5 SDM kit in L0 vectors using primers 15-18 (Supplementary Table 3). L0 Cas9YFP was a gift from Thomas Mock^72^. Level 0 PtU6 promoter was deposited in Addgene (#104895).

Level 1 assembly: FCP promoter, Ble and FCP terminator L0 modules were assembled into L1 pICH47732. FCP promoter, Cas9 and FCP terminator L0 modules were assembled into L1 pICH47742. Level 1 Ble and Cas9 under *P. tricornutum* FCP promoter and terminator were deposited in Addgene (#104893, #104894, respectively). The sgRNA scaffold was amplified from pICH86966_AtU6p_sgRNA_NbPDS^77^ with sgRNA guide sequences integrated through the forward primers 21-23 (Supplementary Table 3). Together with the L0 U6 promoter, sgRNA_1 and sgRNA_2 were assembled into L1 destination vectors pICH47751 and pICH47761, respectively.

Level 2 assembly: L1 modules pICH47732:FCP:Ble, pICH47742:FCP:Cas9YFP, pICH47751:U6:sgRNA_PtMC5 1, pICH47761: U6:sgRNA_PtMC5 2 and the L4E linker pICH41780 were assembled into the L2 destination vector pAGM4723. Constructs were screened by digestion with EcoRV or EcoRI and by PCR. See Supplementary Fig. 6a-c for an overview of the Golden Gate assembly procedure and the final construct.

### Transformations of *P. tricornutum*

Cells were transformed as previously described^78^ using the Bio-Rad Biolistic PDS-1000/He Particle Delivery System fitted with 1550 psi rupture discs. Tungsten particles M17 (1.1 mm diameter) were coated with 5 μg circular plasmid DNA in the presence of 2.5 M CaCl_2_ and 0.1 M spermidine. Approximately 2·10^6^ cells were spread in the center of a plate of a solid medium (50% FSW + f/2,1.5% agar) 2 days before bombardment. For transformation, the plate was positioned at the second level within the Biolistic chamber. Bombarded cells were set to recover for 1 day prior to suspension in 1 ml sterile FSW + f/2. Cell suspension was plated onto solid medium containing 100 μg·ml^-1^ Phleomycin. After 2-3 weeks resistant colonies were re-streaked onto fresh solid medium containing 100 μg·ml^-1^ Zeocin.

### Selection of knockout lines

Resistant colonies were scanned for presence of the Cas9 gene by colony PCR. Cas9 positive colonies were scanned for the size of PtMC5 amplicon (primers 24-25 and 26-27, Supplementary Table 3), colonies exhibiting double-bands, representing both WT (714 bp) and edited (~590 bp) PtMC5 (probably heterozygotes or mosaic colonies) were re-streaked onto fresh solid medium containing 100 μg·ml^-1^ Zeocin. Daughter colonies were scanned for the size of PtMC5 amplicon, colonies exhibiting a single band representing bi-allelic edited PtMC5 (~590 bp) were selected and the PtMC5 gene was sequenced to locate the exact deletion (primers 26-28, Supplementary Table 3).

### PtMC5 expression in *P. tricornutum*

PtMC5 including the 6xHis was ligated into the *P. tricornutum* expression vector, which includes the *P. tricornutum* histone4 promoter, PH4-pPhatT1^33,79^ using EcoRI and KpnI restriction sites, the KpnI restriction site was introduced to the PtMC5 using primers 29-30 (Supplementary Table 3). *P. tricornutum* cells were transformed as above, colonies were scanned for the presence of the expression plasmid by colony PCR using primers 31-32 (Supplementary Table 3). In addition, the presence of the plasmid’s PtMC5 (541 bp) in addition to the endogenous PtMC5, (714 bp) was verified using primers 26-27 (Supplementary Table 3).

### RNA isolation and RT-PCR analysis

RNA was isolated from 50 ml cultures with the Direct-zol RNA miniprep kit (Zymo research) according to the manufacturer’s instructions, followed by DNase treatment with Turbo DNase (Ambion). Equal amounts of RNA were used for cDNA synthesis with the ThermoScript RT-PCR system (Invitrogen). For transcript abundance analysis, Platinum SYBR Green qPCR SuperMix-UDG with ROX (Invitrogen) was used as described by the manufacturer. Reactions were performed on QuantStudio5 Real-Time PCR Systems (ThermoFisher) as follows: 50 °C for 2 min, 95 °C for 2 min, 40 cycles of 95 °C for 15 s, 60 °C for 30 s. The primers for the *PtMC5* gene capture the 1^st^ exon-intron junction and exon 2, detecting WT, OE and KO *PtMC5* (primers 33-34, Supplementary Table 3). Transcript abundance of PtMC5 was calculated by normalizing to expression of TBP^80^ (primers 35-36, Supplementary Table 3) in each sample and to the expression of the WT sample.

### PtMC5 gene and protein modeling

The gene sequence and amino acid sequence of PtMC5 were obtained from the JGI genome portal (protein ID: 54873, transcript ID: estExt_Phatr1_ua_kg.C_chr_160041), and corrected manually using ESTs (the prediction extended the sequences artificially by two exons on the 5’ end which were removed in the final sequence). Conserved domain prediction (CDD https://www.ncbi.nlm.nih.gov/Structure/bwrpsb/bwrpsb.cgi, and Choi^26^) were used to obtain protein domains. Modeling of the 3D structure of PtMC5 was performed online using the SwissModel server (http://swissmodel.expasy.org) based on the structure of *S. cerevisiae* MC (4F6O)^41^ as a template. Molecular graphics were prepared using PyMOL software (http://www.pymol.org/, Schröedinger).

### Identifying MC genes from various species

Initial lists of genes were taken from the pico-Plaza (https://bioinformatics.psb.ugent.be/plaza/versions/pico-plaza/) gene family HOM000388 for the following species: *Aureococcus anophagefferens*, *Arabidopsis thaliana*, *Chlorella sp* NC64A, *Chlamydomonas reinhardtii*, *Coccomyxa sp.* C169, *Ectocarpus siliculosus*, *Fragilariopsis cylindrus*, *Micromonas pusilla* CCMP1545, *Micromonas sp.* RCC299, *Oryza sativa, Physcomitrella patens, Phaeodactylum tricornutum*, *Thalassiosira pseudonana, Volvox carteri*. Sequences were also taken from JGI according to Supplementary Table 1 from Choi^26^, with the exception of *Guillardia theta*, whose sequences were taken from Klemenčič *et al*.^55^, and *Emiliania huxleyi*, where the sequences were manually curated. For the diatoms, sequences were taken from the Moore collection MMETSP44. The MMETSP data is currently available at: http://datacommons.cyverse.org/browse/iplant/home/shared/imicrobe/camera/camera_mmetsp_ncgr/combined_assemblies. All of the diatom datasets were downloaded, both the .pep and the .nt files, for the following species: *Amphiprora sp. Amphora coffeaeformis* CCMP127, *Asterionellopsis glacialis* CCMP134, *Chaetoceros affinis* CCMP159, *Chaetoceros curvisetus*, *Chaetoceros debilis* MM31A_1, *Chaetoceros neogracile* CCMP1317*, Corethron pennatum* L29A3, *Ditylum brightwellii* GSO103, *Ditylum brightwellii* GSO104, *Ditylum brightwellii* GSO105, *Extubocellulus spinifer* CCMP396, *Fragilariopsis kerguelensis* L2_C3, *Fragilariopsis kerguelensis* L26_C5, *Nitzschia punctata* CCMP561, *Proboscia alata* PI_D3, *Pseudonitzschia australis* 10249_10_AB, *Pseudonitzschia fradulenta* WWA7, *Skeletonema dohrnii* SkelB, *Skeletonema marinoi* SkelA, *Skeletonema menzelii* CCMP793, *Thalassionema nitzschioides* L26_B, *Thalassiosira antarctica* CCMP982, *Thalassiosira gravida* GMp14c1, *Thalassiosira miniscula* CCMP1093, *Thalassiosira oceanica* CCMP1005, *Thalassiosira rotula* CCMP3096, *Thalassiosira rotula* GSO102, *Thalassiosira weissflogii* CCMP1010, *Thalassiosira weissflogii* CCMP1336, *Thalassiothrix antarctica* L6_D1. Sequences from *Skeletonema costatum* (Skcos) and an additional isolate of *Thalassiosira rotula* (Throt) were provided by Harriet Alexander and Sonya T. Dyhrman^45^. For species with more than one isolate (*D. brightwellii, F. kerguelensis, T. rotula, T. weissflogii*) the isolate with the most complete MCs was chosen. To identify MCs, a database was built of all the peptide files from the above species, and local Blat was run using the PtMC5 p20 domain as the query. An additional database was built of the nucleotide sequences, and a translated Blat was run to identify additional potential transcripts that were not translated. For each isolate, manual curation was performed to eliminate redundancy. Sequences from *Hemiaulus sinensis* were found by Blast search at NCBI in the TSA database, and *Fistulifera solaris* were found by Blast search at NCBI in the protein database. For non-diatom species, the input file for database searches was of the full length MCs of TpMC1-6, PtMC5 and SmMC1-5, and the searches run were translated Blasts (using the protein query against the species DNA). Red algae: The following genomes were searched at Ensembl Plants: *Chondrus crispus* (hits), *Cyanidioschyzon merolae* (none), *Galdieria sulphuraria* (none), also checked genome, local. For *Porphyridium purpureum* genome http://cyanophora.rutgers.edu/porphyridium/ website was used. *Porphyra yezoensis* (Nori) search performed at NCBI (genome, TSA). *Nannochloropsis gaditiana* (genome local, TSA at NCBI), *Cyanophora paradoxa* local searches. *Symbodinium* isolates: A1, A2, B2, C, D, *muscatinei*: Translated Blast was run at NCBI against the TSA database. For each isolate manual curation was performed to obtain a non-redundant dataset. A1 was chosen as it had the most and most complete MCs. *Plasmodium falciparum* was taken as a representative of *Plasmodium* species^81^. All the full protein sequences of the MCs are presented in Supplementary Data 1.

### Identifying p20 and p10 domains

The putative protein sequences of the various MCs were run against the CDD database at NCBI to find the p20 domain. For many of the sequences the definition of the p10 domain as available in the public domain databases (Pfam, CDD, InterPro) did not find hits. Based on alignment of our sequences and the supplemental alignment^26^, we built new patterns to search for the p10 in the various diatom sequences. The basis of the p10 pattern were a sequence of [QE]TSAD at the beginning and GAX[ST]XXXXXX[IVLA] in the middle. This was refined to Dx[QE]TSAD at the beginning, GGAX[ST] in the middle and QxPQL at the end of the putative p10. The patterns were the basis for the search which was performed manually on all of the defined diatom MCs. The putative p10 and p20 domains are marked on the full MCs protein sequences in Supplementary Data 1.

### Phylogenetic tree preparation

Alignments were performed on whole protein sequences, on the p20 and p10 domains. The p20 domains were trimmed manually from the full length sequences based on alignment to the CDD database, and further refined by hand. The p10 domains were aligned as described (in the other file). Alignments were performed using ClustalW 2.1. Phylogenetic trees were built using the Neighbor-joining algorithm in ClustalW and with Maximum likelihood (ProML) in the Phylip package. Trees were visualized using the iTol server (https://itol.embl.de/).

### Statistical analysis

All reported *P*-values were determined using a two-tailed unpaired Student’s t-test. In all figures, error bars represent s.e. and ‘n’ defined as the number of unrelated replicas of each treatment. Box plots were produced using BoxPlotR web tool (http://shiny.chemgrid.org/boxplotr/).

## Acknowledgments

We are grateful to Harriet Alexander and Sonya T. Dyhrman for assistance with metatranscriptome data sets from Narragansett Bay. We thank Yishai Levin for the help with analysis of the redox proteomics. We thanks Adi Volpert and Inbal Nussbaum for technical assistance. We thanks Daniella Schatz for valuable feedback

This research was supported by the Israeli Science Foundation (ISF) (grant # 712233) awarded to AV.

